# A bone marrow stromal secretome screen identifies semaphorin 3A as a regulator of hematopoiesis

**DOI:** 10.1101/2025.08.07.669162

**Authors:** Daniel K Borger, Shane PC Mitchell, Fumio Nakahara, Daniele Filippo Colombo, Lidiane S Torres, Scott Garforth, Simone Sidoli, Jeroen Krijgsveld, Lev Silberstein, Paul S Frenette, Kira Gritsman

**Affiliations:** Ruth L. and David S. Gottesman Institute for Stem Cell and Regenerative Medicine Research, Albert Einstein College of Medicine, Bronx, NY, USA; Department of Cell Biology, Albert Einstein College of Medicine, Bronx, NY, USA; Division of Regenerative Medicine, Center for Molecular Medicine, Jichi Medical University, Shimotsuke, Tochigi, Japan; Division of Infectious Diseases, Advanced Clinical Research Center, Institute of Medical Science, The University of Tokyo, Minato-ku, Tokyo, Japan; Division of Proteomics of Stem Cells and Cancer, German Cancer Research Center (DKFZ), Heidelberg, Germany; Department of Biochemistry, Albert Einstein College of Medicine, Bronx, NY, USA; Medical Faculty, Heidelberg University, Heidelberg, Germany; Fred Hutchinson Cancer Center, Seattle, WA, USA; Department of Medicine, Albert Einstein College of Medicine, Bronx, NY, USA; Department of Medical Oncology, Albert Einstein College of Medicine, Bronx, NY, USA

## Abstract

Bone marrow mesenchymal stromal cells (MSCs) are a major source of secreted factors that control hematopoietic stem and progenitor cell (HSPC) function. We previously reported the generation of revitalized MSCs (rMSCs), which more effectively support HSPCs in culture. In a secretome screen using rMSCs, we identified semaphorin 3A (SEM3A) as a secreted factor upregulated as part of a pro-inflammatory signature that may contribute to HSPC expansion by rMSCs. We show that recombinant SEM3A acts directly on HSPCs to inhibit their cycling *ex vivo*. Analysis of a SEM3A loss of function mutation *in vivo* revealed hematopoietic progenitor expansion and accelerated recovery after myeloablation, consistent with a role for SEM3A in regulating HSPCs at steady state and during hematopoietic stress. This work highlights proteomic screening using rMSCs as a method to identify novel secreted niche factors and uncovers a novel role for SEM3A in controlling HSPC proliferation in stress hematopoiesis.

**Summary:** Borger et al. characterize the secretome of revitalized bone marrow stromal cells and identify a novel role of the protein semaphorin 3A in regulating hematopoietic stem and progenitor cell proliferation in steady state and stress conditions.

## Introduction

Hematopoietic stem and progenitor cells (HSPCs) underpin the entirety of the adult hematopoietic system, the source from which all cells of the blood and immune system are continuously replenished. To achieve this throughout the lifetime of an organism, HSPCs must be preserved through quiescence and self-renewal, differentiate into the necessary lineages at the appropriate rates, and be able to respond to hematopoietic emergencies, such as infection or blood loss (Kasbekar et al., 2023). While HSPCs possess the ability to recognize and respond directly to some environmental cues such as pathogens (Demel et al., 2022; Takizawa et al., 2011), many of these behaviors are regulated locally in the bone marrow by a dense network of cellular and non-cellular support, collectively termed the HSPC niche. Niche cells support HSPCs and coordinate their behavior through paracrine and juxtacrine signals (Pinho and Frenette, 2019). Although these niche signals have been a major focus of research, significant gaps in our knowledge remain, especially with regard to how such signals are modified in the setting of hematopoietic emergencies.

In response to bleeding (Cheshier et al., 2007), infection (Baldridge et al., 2010), or myeloablation (Wilson et al., 2008), HSCs will be activated transiently, entering the cell cycle to replenish more committed progenitors and, in some cases, to rapidly differentiate into mature cells (Haas et al., 2015). While HSCs that cycle in response to inflammatory challenges lose stem cell function to some degree (Bogeska et al., 2022), re-entry into a quiescent state is thought to be essential for preserving stem cell function and preventing rapid HSC depletion in response to inflammation (Pietras et al., 2014; Wilson et al., 2008). In addition to endogenous brakes on cell cycling (Chavez et al., 2021; Traveset et al., 2024), niche-derived signals such as TGFβ (Herault et al., 2017; Zhao et al., 2014) have been implicated in this re-enforcement of HSC quiescence. Additionally, evidence suggests that the response to hematopoietic stress is heterogeneous among HSCs, with small numbers of HSCs preserved in a quiescent state even as the bulk of the HSC pool is activated (Bogeska et al., 2022; Wilson et al., 2008; Zhao et al., 2019). However, the precise mechanisms and signaling factors underlying the maintenance of quiescent HSC subsets during stress hematopoiesis and the return to quiescence by cycling HSCs after its resolution remain unclear.

Mesenchymal stromal cells (MSCs) are a critical component of the HSPC niche, expressing factors such as stem cell factor (SCF) and C-X-C motif chemokine ligand 12 (CXCL12), which are indispensable for HSPC maintenance and retention in the bone marrow (Asada et al., 2017; Barker, 1994; Greenbaum et al., 2013; Sugiyama et al., 2006). Proteomic characterization of the MSC secretome could be useful for identifying additional factors underlying niche interactions, but these cells show limited ability to support HSPCs *ex vivo*, due in part to downregulation of SCF and CXCL12 (Nakahara et al., 2019). This makes using primary MSCs for *ex vivo* secretomic analysis challenging. Previously, we identified five factors – *Klf7*, *Ostf1*, *Xbp1*, *Irf3*, and *Irf7*, collectively termed the KOXII factors – that, after transduction into primary murine MSCs, increased expression of secreted and surface proteins critical for HSPC support. These rMSCs could expand functional HSCs in *ex vivo* culture settings without supplemental cytokines, unlike empty-vector transduced control MSCs (cMSCs) (Nakahara et al., 2019).

In the present study, we characterize the secretome of these rMSCs in an effort to better understand how these factors drive this revitalization of niche function and to identify proteins with unrecognized roles in niche-HSPC crosstalk. We find that rMSCs exhibit features of MSCs in inflammatory settings, granting some insight into how these cells drive HSPC expansion *ex vivo*. Additionally, we find that semaphorin 3A (SEM3A), a secreted factor, is highly upregulated in the secretome of rMSCs. SEM3A was initially identified for its role in axonal guidance via its binding to class A plexins and their co-receptor neuropilin-1 (NRP1) (Sharma et al., 2012). NRP1-dependent roles for semaphorin 3a in direct regulation of bone marrow vascular regeneration and bone remodeling have been identified more recently (Hayashi et al., 2012; Termini et al., 2021). However, to date, there is no evidence of direct regulation of HSPCs by SEM3A. Here, we demonstrate that SEM3A promotes HSC quiescence directly, and that loss of SEM3A signaling *in vivo* leads to an increase in HSPC numbers, mediated at least in part by de-repression of cytokine signaling. Additionally, we show that SEM3A is upregulated by MSCs *in vivo* in response to hematopoietic stress, suggesting a role in controlling HSPC proliferation in inflammatory settings. Altogether, this work provides a new secretomic dataset that will be useful in understanding interactions between the MSCs and HSPCs, especially in inflammatory settings. Furthermore, it provides evidence for a novel role for SEM3A in direct regulation of HSPC proliferation.

## Results

### The rMSC transcriptome exhibits inflammatory signatures

We previously reported that rMSCs can secrete increased levels of niche factors such as stem cell factor (SCF) and support *ex vivo* expansion of engraftable HSCs (Nakahara et al., 2019). Two of the KOXII factors, *Irf3* and *Irf7*, are critical factors both in the induction of type I interferon expression in response to pathogen-associated molecular patterns and in the cellular response to interferon signaling (Honda and Taniguchi, 2006). Both *Irf3* and *Irf7* are indispensable for the ability of the KOXII factors to drive HSC expansion by rMSCs *ex vivo* (Nakahara et al., 2019). This suggests that the *ex vivo* HSC expansion achieved by rMSCs may be in part due to rMSCs mimicking *in vivo* MSC inflammatory responses. Consistent with this hypothesis, gene set enrichment analysis (GSEA) of RNAseq data from rMSCs compared with empty-vector transduced control MSCs (cMSCs) revealed that rMSCs exhibited signatures of interferon α (IFNα) signaling, as well as an MSC-specific poly-I:C response signature derived from previously published data (Helbling et al., 2019; Subramanian et al., 2005) (**Fig. 1 A**). As type I interferon signaling is critical for driving HSC activation in certain infections and in response to myeloablation (Clapes et al., 2021; Essers et al., 2009; Traveset et al., 2024), we hypothesized that rMSCs can be used as an *ex vivo* model system to study the factors important for controlling HSC cycling and self-renewal in the context of hematopoietic stress.

**Figure 1.**
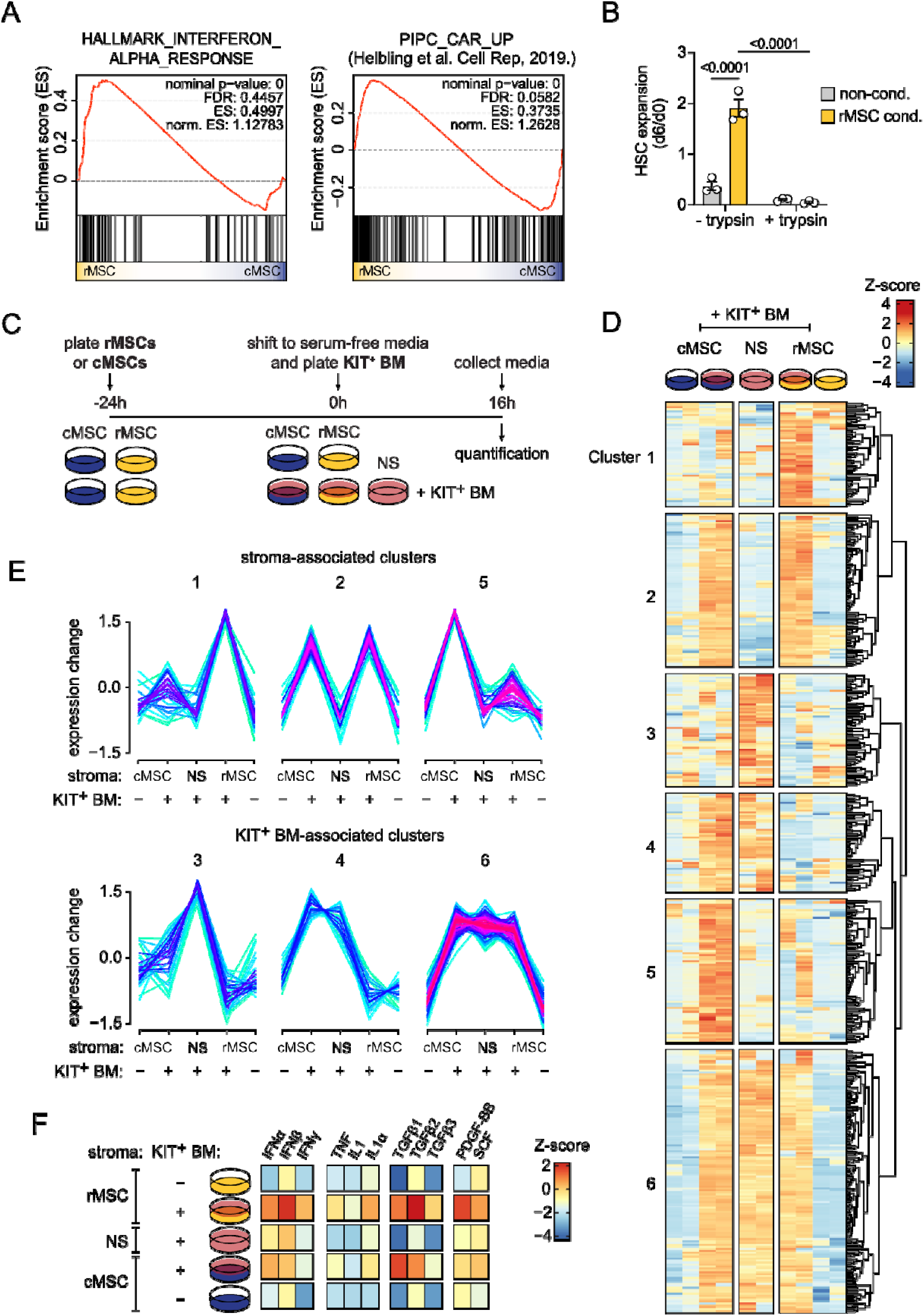
Proteomic characterization of the rMSC secretome. (**A**) GSEA performed on RNA-seq of rMSCs and cMSCs, showing enrichment of two gene sets associated with interferon signaling within rMSCs. PIPC_CAR_UP is a gene set consisting of the most highly upregulated genes (> 2 log fold change) in PDGFRβ^+^ Sca-1^-^ murine bone marrow MSCs after PIPC injection (Helbling et al., 2019). (**B**) Lin-depleted BMNCs were cultured in non-conditioned or rMSC conditioned media with (+ trypsin) or without (- trypsin) pretreatment with trypsin followed by trypsin inhibitor. Expansion of phenotypic HSCs (CD150^+^ CD48^-^ LSKs) was determined by flow cytometry. ANOVA with Tukey’s multiple comparison test was used to compare conditions. (**C**) Approach for unlabeled proteomic characterization of the rMSC and cMSC secretome in both monoculture and co-culture with KIT-enriched hematopoietic cells (KIT^+^ BM). (**D**) Heatmap of proteomic results. Proteins were clustered using fuzzy C means clustering algorithm based on patterns of detection (Schwammle and Jensen, 2010). Proteins with cluster membership degree greater than 0.4 are shown. (**E**) Pattern of protein detection of all fuzzy C means clusters shown in (**D**). Clusters of proteins detected in culture of KIT^+^ BMNCs alone were termed KIT^+^-associated clusters, while those appearing only in stromal culture conditions were termed stroma-associated clusters. (**F**) Selected results of IPA upstream regulator analysis (URA) performed on proteomics hits.

### Proteomics reveals that SEM3A is elevated in the rMSC secretome

To determine the degree to which secreted factors play a role in the expansion of HSCs by rMSCs, lineage-depleted (Lin^-^) bone marrow mononuclear cells (BMNCs) were cultured in conditioned media collected from rMSCs. After 6 days in culture with conditioned media, phenotypic Lin^-^ SCA-1^+^ KIT^+^ (LSK) CD48^-^ CD150^+^ HSCs numbers were determined by flow cytometry. rMSC conditioned media expanded HSCs approximately two-fold over input, whereas culture in non-conditioned media led to a decline in HSC numbers (**Fig. 1 B**). As expected, pre-treatment with trypsin completely abrogated the ability of rMSC conditioned media to expand HSCs, suggesting that this effect is primarily mediated by secreted proteins (**Fig. 1 B**).

To identify the proteins that mediate HSC expansion by rMSC conditioned media, we performed unlabeled proteomics on cell-free media supernatants from rMSCs and cMSCs cultured alone (no stroma, NS) or in co-culture with KIT-enriched BMNCs (KIT^+^ BM) (**Fig. 1 C**). This approach detected 446 proteins across all conditions (**Table S1**). Interestingly, we saw that hematopoietic co-culture led to broad increases in protein secretion in both cMSC and rMSC conditions, suggesting that hematopoietic cell–MSC crosstalk leads to major changes in the secretory behavior of MSCs, or that the presence of hematopoietic cells drives large scale release of factors previously trapped within the ECM (**Fig. 1 D**). Fuzzy c-means clustering (Schwammle and Jensen, 2010) of these proteomic hits yielded six groups of proteins with differing patterns of detection across the different culture settings. Most notably, three clusters demonstrated enrichment in the co-culture settings, with cluster 1 enriched in the rMSC secretome in co-culture, cluster 5 enriched in the cMSC secretome in co-culture, and cluster 2 showing similar enrichment in both (**Fig. 1, D and E**). STRING graphing (Szklarczyk et al., 2025) of proteins found in cluster 5, which is enriched in the cMSC secretome, demonstrated large groups of cytoskeletal components and protein synthesis machinery, suggestive of increased cell death in the cMSC setting (**Fig. S1, A and B**). Within cluster 1, which was enriched in the rMSC secretome in hematopoietic co-culture, we saw a number of extracellular matrix (ECM) components, ECM-modifying enzymes, and ECM-binding cell surface proteins (**Fig. 1 E and Fig. S1 C**). Upstream Regulator Analysis (URA), a function of Ingenuity Pathway Analysis (Qiagen), revealed signatures of a number of pro-inflammatory cytokines which were enriched in the rMSC co-culture settings, most notably three members of the interferon family, IFNα, IFNβ, and IFNγ (**Fig. 1 F**). This is in keeping with the IFN signatures found in the RNAseq data, supporting the view that rMSCs mimic an inflamed niche (**Fig. 1 A**).

Among the proteins in cluster 1, consisting of proteins upregulated in the rMSC secretome in hematopoietic co-culture, was semaphorin 3A (SEM3A, encoded by *Sema3a*), a soluble secreted signaling factor (**Fig. 2 A**). When comparing the rMSC and cMSC secretomes in co-culture settings, SEM3A was significantly upregulated in the rMSC secretome, corresponding to its upregulation in rMSCs at the RNA level (**Fig. 2 B and Fig. S1 D**). The increase in SEM3A protein secretion in rMSC conditioned media was confirmed by ELISA (**Fig. 2 C**). As our co-culture proteomic approach cannot directly distinguish between MSC-derived and hematopoietic cell-derived proteins, we performed qPCR on sorted stromal and hematopoietic cells after co-culture. As expected, *Sema3a* was expressed by rMSCs but not hematopoietic cells, identifying the former as a predominant source of SEM3A in the co-culture setting (**Fig. 2 D**).

**Figure 2.**
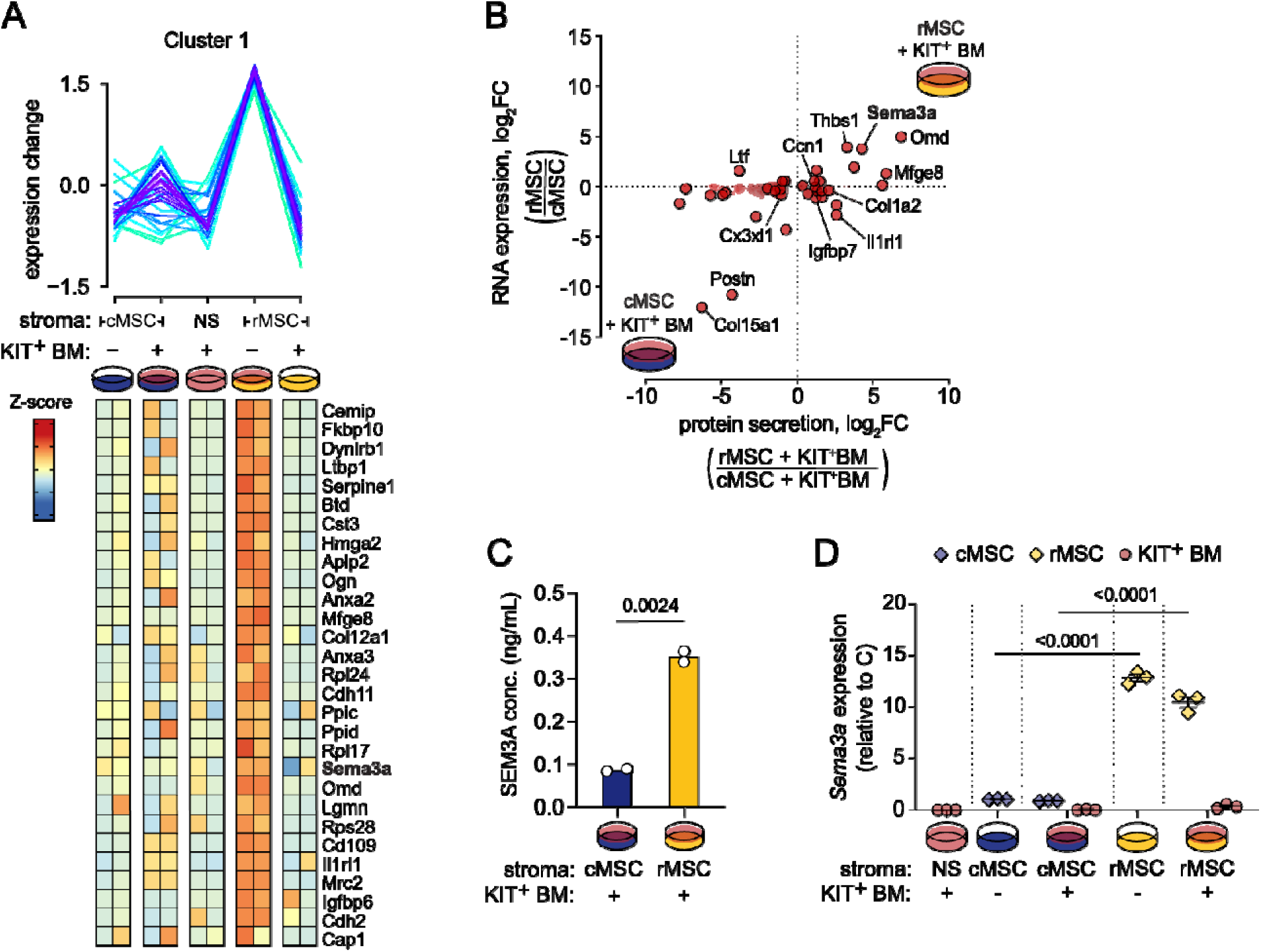
SEM3A is upregulated in the rMSC secretome. (**A**) Pattern of protein detection of cluster 1, demonstrating increased levels in the rMSC secretome in co-culture conditions. Heatmap shows proteins assigned to cluster 1 with membership degree greater than 0.4. (**B**) log_2_FC of normalized LFQ of proteins in rMSC secretome versus cMSC secretome in co-culture conditions, plotted against log_2_FC of RPKM of corresponding transcripts, as determined by RNAseq. Significantly differentially detected proteins (p < 0.05) are indicated in red, and extracellular proteins per STRING indicated in dark red. (**C**) Results of SEM3A ELISA performed on cMSC and rMSC conditioned media. Unpaired two-tailed Student’s *t* test was used to compare conditions. (**D**) *Sema3a* expression, as determined by qPCR, in FACS-purified hematopoietic cells (KIT^+^), cMSCs, or rMSCs after being subjected to the same conditions indicated in **Fig. 2 A**, including either monoculture or co-culture. ANOVA with Tukey’s multiple comparison test was used to compare conditions.

To confirm the secretomic hits from label-free proteomics, we also performed pulsed stable isotope labelling of amino acids in culture (pSILAC) on rMSCs and cMSCs (**Fig. S2 A**). pSILAC detected a similar number of total proteins as the label-free approach (510 proteins across both conditions). Of these, 141 were isotope-labelled and quantifiable across conditions. SILAC ratios could be calculated for 72 proteins, demonstrating upregulation of several extracellular matrix (ECM) and osteoblast-related proteins (**Fig. S2 B and Table S2**). We also compared label-free quantification (LFQ) intensity of all 141 labelled proteins after imputing missing intensities for proteins detected in at least one replicate (**Fig. S2 C and D; Table S3**). STRING graphing of proteins upregulated in the rMSC secretome again revealed a cluster of ECM-related proteins, including several proteins previously seen upregulated in the rMSC secretome by label-free proteomics (**Fig. S2 E**). Importantly, pSILAC confirmed that SEM3A was upregulated in the rMSC secretome (**Fig. S2 C and D**).

### SEM3A directly regulates HSPC proliferation *ex vivo*

SEM3A has previously been identified as an important factor in controlling osteoblast differentiation and vascular regeneration in the bone marrow, but a direct role in regulating HSPCs has not been explored (Hayashi et al., 2012; Termini et al., 2021). To address this question, FACS-purified mouse bone marrow LSK (Lin-SCA-1+ KIT+) cells were cultured in PVA-based expansion media (Wilkinson et al., 2019; Wilkinson et al., 2020) in the presence of either a recombinant SEM3A-Fc fusion protein or an IgG2a isotype control antibody (10μg/mL) for 4 days. At day 4, total cell numbers were moderately increased (1.18-fold) in SEM3A-Fc-treated cultures relative to IgG2a (**Fig. 3 A**). LSK numbers were similarly increased in SEM3A-Fc treated cultures, although LSK frequency was not significantly increased (**Fig. 3, B and C**). Notably, compared to IgG2a-treated cultures, SEM3A-Fc treated cultures demonstrated lower overall levels of cell death, which may explain this broad expansion in cell numbers (**Fig. S3 A**). To determine if SEM3A has a specific effect on HSCs, we assessed the numbers of CD150+ EPCR+ LSK cells, as EPCR expression has been shown to better correspond to HSC function in these culture settings (Che et al., 2022; Zhang et al., 2012). In contrast to what was observed with LSKs, we saw an increase in HSC frequency with SEM3A-Fc treatment, translating to a 1.44-fold increase in HSC numbers (**Fig. 3, D and E**). This was abrogated by addition of a SEM3A neutralizing antibody, confirming that the observed HSC expansion is driven by SEM3A-Fc rather than a result of endotoxin or other contaminants (**Fig. S3 B**).

**Figure 3.**
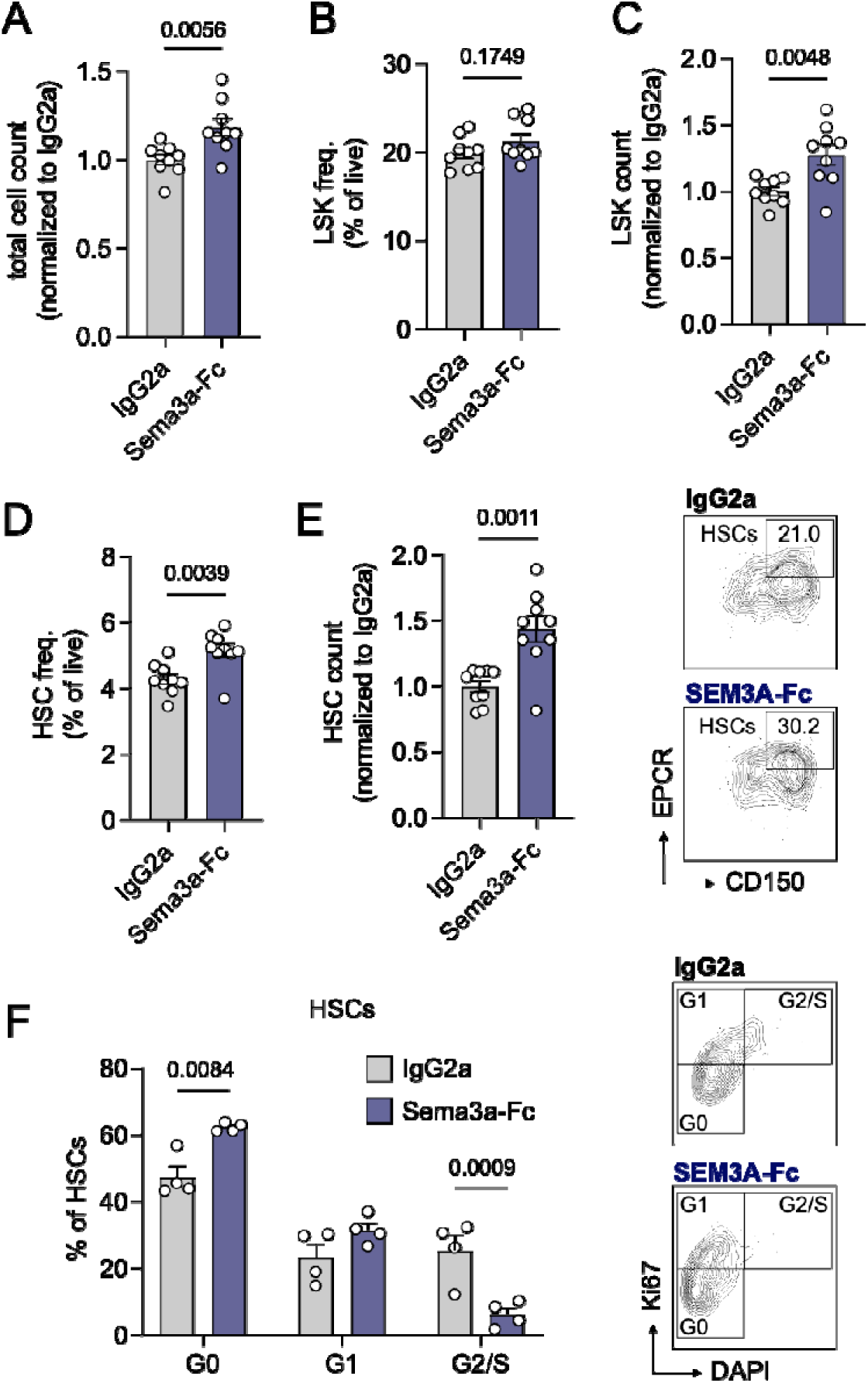
SEM3A preserves HSPCs *ex vivo* by regulating proliferation. (**A**) Total cell count, (**B**) LSK frequency and (**C**) count, and (**D**) CD135-CD150+ EPCR+ LSK (HSC) frequency and (**E**) count after 4d of culture of 2×10^3^ LSKs with recombinant SEM3A-Fc or IgG2a isotype control. Representative flow plots gated on CD135^-^ LSKs are shown. Results shown are from cells derived from three animals and two independent experiments. Unpaired two-tailed Student’s *t* test was used to compare conditions. (**F**) Cell cycle analysis of CD150+ EPCR+ HSCs after 48h of culture of 1×10^4^ LSKs. Representative DAPI/Ki-67 plots, gated on HSCs, are shown. This experiment was performed three times, with similar results. ANOVA with Tukey multiple-comparisons test was used to compare conditions.

To investigate whether SEM3A-Fc influences HSC expansion by modulating cell cycle status, we performed cell cycle analysis using DAPI and Ki-67 staining on LSKs cultured with SEM3A-Fc or IgG2a isotype for 48h. As expected for these culture settings, we observed much higher levels of HSC cycling than is typically seen at steady state *in vivo* (**Fig. 3 F**). Interestingly, SEM3A-Fc promoted HSC quiescence, with HSCs in the SEM3A-Fc treated conditions showing an increase in G0 occupancy and a decrease in late cell cycle (G2/S) (**Fig. 3 F**). Together, these data suggest that, rather than drive HSC expansion by promoting entry into the cell cycle, SEM3A increases HSC numbers *ex vivo* by promoting HSC quiescence and survival, thereby preserving HSC identity in a setting of high HSC cycling.

### *Sema3a* is expressed in the HSPC niche

The effects that SEM3A exerted on HSPC cycling *ex vivo* suggests a possible HSPC niche function for SEM3A. To assess whether SEM3A plays a role in the HSPC niche *in vivo*, we first assessed *Sema3a* expression across the major cellular compartments of the bone marrow in healthy adult mice. qPCR for *Sema3a* for stromal populations, HSPCs, and mature immune cells demonstrated high SEM3A expression in sinusoidal endothelial cells (SECs), with limited expression in MSCs (**Fig. 4 A and Fig. S4 A**). Expression of *Sema3a* in HSPCs and mature immune cells was low or undetectable compared with expression in these niche populations, confirming that SEM3A expression is limited to the bone marrow stroma (**Fig. 4 A**). Analysis of a compilation of existing steady state murine bone marrow stroma single cell RNA sequencing (scRNAseq) datasets confirmed this pattern of expression (Baccin et al., 2020; Baryawno et al., 2019; Dolgalev and Tikhonova, 2021; Tikhonova et al., 2019). Of the major cell populations associated with HSPC niche function within these datasets – broadly, sinusoidal endothelial cells (SECs), arteriolar endothelial cells (AECs), and MSCs – *Sema3a* expression was largely restricted to the SEC compartment (**Fig. S4 B**). Examining the expression pattern of all semaphorin family member in bulk RNAseq of bone marrow endothelial cells and MSCs confirmed that *Sema3a* belongs to a subset of semaphorins predominantly expressed by endothelial cells (**Fig. S4 C**) (Asada et al., 2017). This is in line with previous studies that have identified a major autocrine role for SEM3A in regulating vascular remodeling and regeneration (Guttmann-Raviv et al., 2007; Serini et al., 2003; Termini et al., 2021).

**Figure 4.**
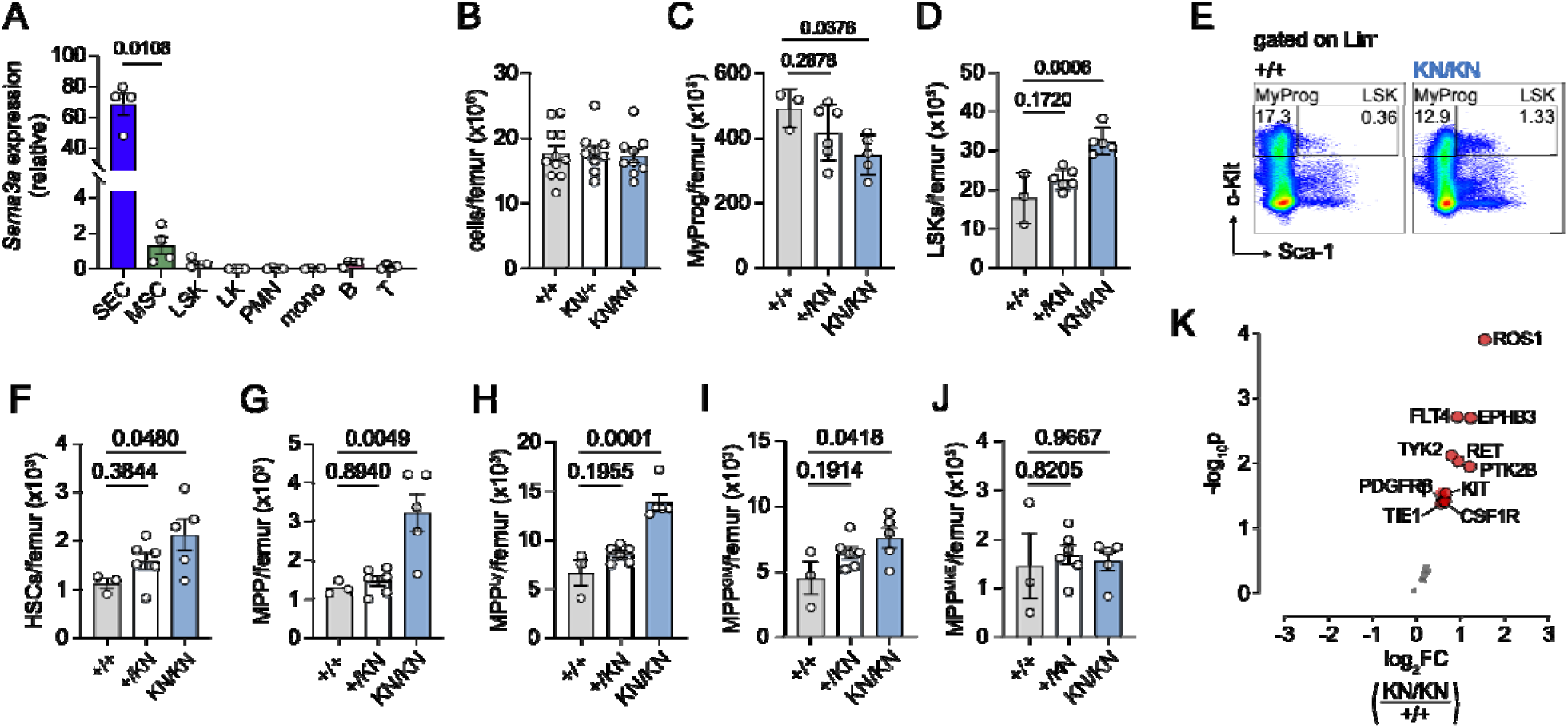
Disruption of SEM3A signaling leads to HSPC expansion *in vivo*. (**A**) Expression of *Sema3a* in CD31+ P-selectin+ Sca-1+ SECs, CD51+ PDGFRa+ MSCs, myeloid progenitors (MyProg), LSKs, neutrophils (PMN), monocytes (mono), B cells, and T cells, as determined by qPCR. Expression normalized to GAPDH. ANOVA with Tukey multiple-comparisons test was used to compare across populations. (**B**) bone marrow cellularity, (**C**) myeloid progenitor (MyProg) numbers, and (**D**) LSK numbers per femur in *Sema3a*^+/+^, *Sema3a*^+/K108N^, and *Sema3a*^K108N/K108N^ mice. (**E**) Representative flow plots gated on Lin^-^ cells corresponding to data shown in **B** and **C**. (**F**) HSC, (**J**) MPP (**G**) MPP^Ly,^ (**H**) MPP^GM^, (**I**) and MPP^MkE^ numbers per femur of *Sema3a*^K108N^ mice. For **B** - **D** and **F** - **J**, ANOVA with Tukey multiple-comparisons test was used to compare groups (n = 3-6 males per group). (**K**) KIT-enriched cells were isolated from *Sema3a*^+/+^ and *Sema3a*^KN/KN^ mice and phosphorylation of 71 RTKs was determined via phospho-protein array. Volcano plot shows log_2_FC of detection of phosphorylated proteins. Significantly differentially detected proteins (p < 0.05) are indicated in red, with selected proteins outlined and labelled (n = 2).

### *Sema3a*^K108N^ mutation leads to HSPC expansion *in vivo*

To explore the potential role of SEM3A in controlling HSPC behavior *in vivo*, we analyzed mice bearing a germline *Sema3a*^K108N^ loss-of-function mutation (Merte et al., 2010). The K108 residue mutated in these mice resides within the plexin binding surface of the semaphorin domain, and the K108N mutation is thought to disrupt binding to class A plexins and downstream signaling without affecting SEM3A homodimerization or NRP1 co-receptor binding (Janssen et al., 2012; Merte et al., 2010). Notably, analysis of existing RNAseq data shows that several of the class A plexins are expressed in HSCs, suggesting that targeting interactions with class A plexins would interfere with SEM3A signaling in these cells (**Fig. S4 D and E**) (Cabezas-Wallscheid et al., 2017; Cabezas-Wallscheid et al., 2014). *Sema3a*^KN^ homozygotes are also viable and fertile, while exhibiting many of the neurological features associated with other *Sema3a* loss-of-function mutations (Behar et al., 1996; Merte et al., 2010; Taniguchi et al., 1997).

At steady state, *Sema3a*^KN/+^ heterozygotes and *Sema3a*^KN/KN^ homozygotes both exhibited mildly reduced body weight compared to *Sema3a*^+/+^ homozygote wildtype mice but did not show any changes in peripheral blood counts, spleen size, or overall bone marrow cellularity (**Fig. 4 B and Fig. S5 A–E**). However, *Sema3a*^KN/KN^ mice exhibited a 1.8-fold expansion of the LSK compartment compared to *Sema3a*^+/+^ littermate controls (**Fig. 4, D and E**). This LSK expansion occurred at the expense of myeloid progenitors (MyPro), which declined in number in the *Sema3a*^K108N/K108N^ mice (**Fig. 4, C and E**). While *Sema3a*^KN/+^ heterozygotes exhibited intermediate MyPro and LSK numbers, they did not significantly differ from *Sema3a*^WT^ homozygotes (**Fig. 4, C and D**). Within the LSK compartment, HSCs (1.9-fold increase), multipotent progenitors (MPPs, 2.5-fold increase), and lymphoid-biased MPP^Ly^ (2.1-fold increase) showed larger increases compared to myeloid-biased MPPs (MPP^GM^ and MPP^MkE^, 1.7 and 1.2-fold increases, respectively) (Challen et al., 2021) (**Fig. 4, F-J**). This data is consistent with SEM3A acting as a negative regulator of HSPC numbers *in vivo*.

To better understand how the *Sema3a*^KN^ mutation alters signaling in murine HSPCs, we evaluated phosphorylation of 71 receptor-tyrosine kinases in KIT-enriched cells from both *Sema3a*^+/+^ controls and *Sema3a*^KN/KN^ mice using a phospho-protein array (Mouse RTK Phosphorylation Array C1; RayBiotech). Strikingly, almost all of the assayed proteins showed some degree of increased phosphorylation in *Sema3a*^KN/KN^ mice, with 13 proteins demonstrating significant hyperphosphorylation, and no proteins showing a significant decrease in phosphorylation (**Fig. 4 K**). Notable among the proteins with increased phosphorylation was KIT, the receptor for SCF. To determine if this increase in KIT phosphorylation was due to an increase in SCF expression in niche cells, we evaluated *Kitl* RNA expression in both MSCs and SECs, the major sources of SCF in the bone marrow. No changes in *Kitl* expression were detectable in either of these cell types in *Sema3a*^+/+^ and *Sema3a*^KN/KN^ mice. This suggests that the increase in KIT phosphorylation is a downstream effect of reduced *SEM3A* signaling in HSPCs, rather than an effect of altered stromal expression of *Kitl* (**Fig. S5, G and H**). Additionally, SEM3A protein levels are not significantly elevated in *Sema3a*^KN/KN^ mice as measured by ELISA, ruling out gain-of-function effects due to compensatory overexpression of the mutant protein (**Fig. S5 I**). Overall, these data support the view that SEM3A acts to restrict HSPC proliferation by dampening growth factor signaling, with the *Sema3a*^K108N^ mutation leading to relaxation of these restrictions and subsequent expansion of the progenitor pool.

### *Sema3a*^K108N^ mutation accelerates myeloid recovery after 5-fluorouracil

The relative paucity of *Sem3a* expression in MSCs at steady state *in vivo* was intriguing, considering that SEM3A was detected at high levels in the secretome of rMSCs, which exhibit signatures of inflammatory signaling. Hence, we hypothesized that inflammatory signals may drive expression of *Sema3a* in bone marrow MSCs. Inflammatory cytokine signaling is critical to hematopoietic recovery after bone marrow ablation, such as that achieved by the chemotherapeutic 5-fluorouracil (5-FU) (Hemmati et al., 2019; Pietras et al., 2016). To evaluate if bone marrow ablation and subsequent changes in inflammatory signaling within the bone marrow changed patterns of *Sema3a* expression, we treated wild-type mice with 5-fluorouracil (5-FU), sorted bone marrow MSCs and SECs at day 7 post-5-FU, when bone marrow cellularity has reached its nadir, and performed qPCR for *Sema3a* (**Fig. 5 A**). Indeed, while we did not detect changes in *Sema3a* expression within the SEC compartment (**Fig. 5 B**), we detected a significant 3-fold upregulation of *Sema3a* expression in MSCs at d7 post-5-FU (**Fig. 5 C**), supporting the hypothesis that *Sema3a* is a component of the MSC response to inflammation and hematopoietic stress. To determine if SEM3A plays a functional role in the bone marrow during hematopoietic stress, we injected *Sema3a*^+/+^ and *Sema3a*^KN/KN^ mice with 5-FU and evaluated subsequent peripheral blood recovery. *Sema3a*^KN/KN^ mice demonstrated mildly accelerated erythrocyte (RBC) recovery and significantly higher platelet counts at d14 following 5-FU (**Fig 5, D-F**; **Fig. S5**, **J**-**L**). Together, these data support a role for SEM3A in regulating the response of HSPCs to hematopoietic stress.

**Figure 5.**
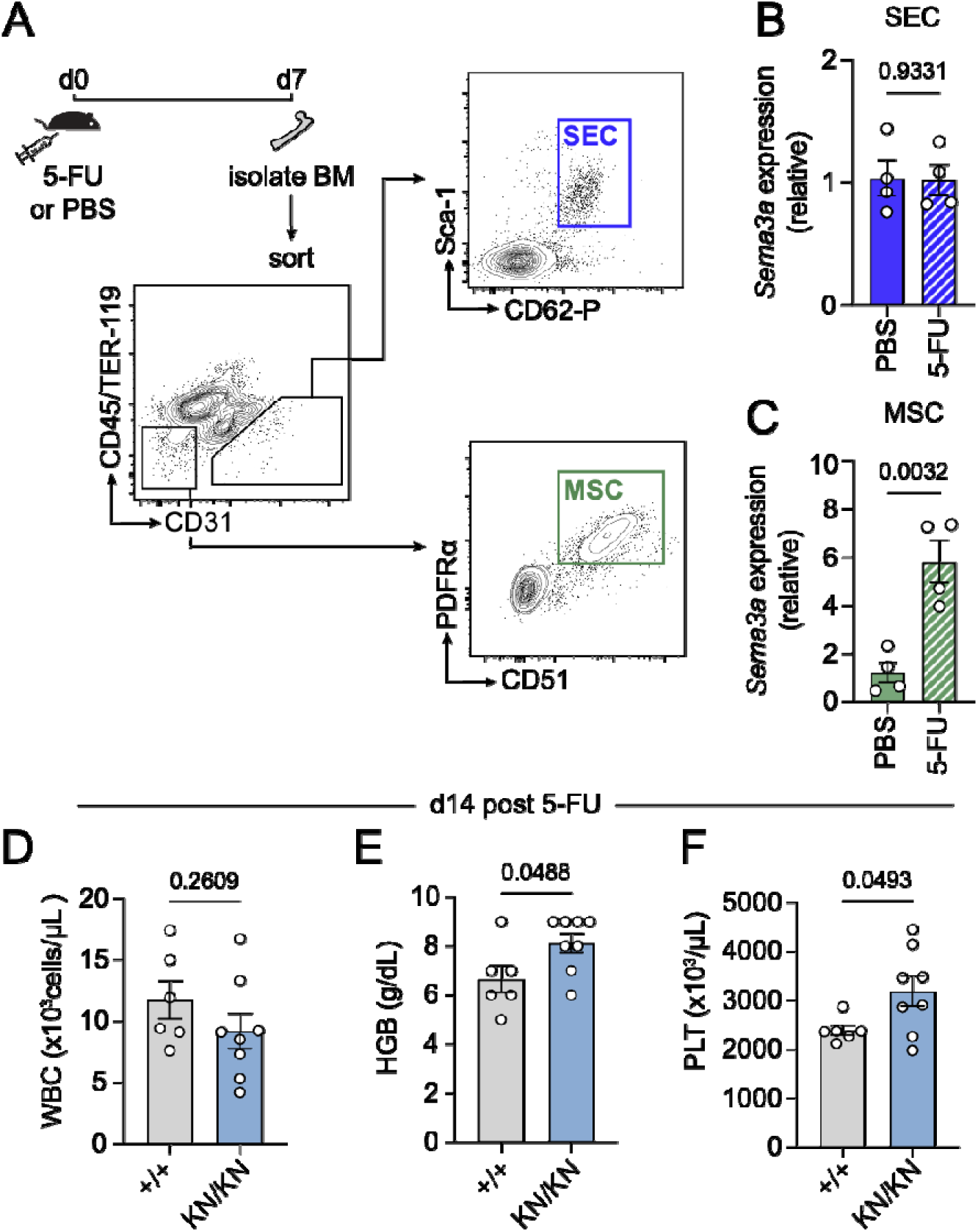
SEM3A mutation alters dynamics of hematopoietic recovery. (**A**) Experimental approach to evaluating expression of *Sema3a* in the HSPC niche following 5-FU treatment. (**B**) Expression of *Sema3a* by SECs and (**C**) MSCs 7d after injection of 5-FU treatment. (**D**) Total white blood cell counts (WBC), (**E**) hemoglobin concentration (HGB), and (**F**) platelet counts (Plts) in peripheral blood of *Sema3a*^+/+^ and *Sema3a*^KN/KN^ mice 14 days after 5-FU injection. For **B** - **F**, unpaired two-tailed Student’s *t* test was used.

## Discussion

While a great deal of effort has been directed towards better understanding the extrinsic signals involved in regulating HSPC behavior, there have been few studies that have leveraged a proteomic approach in identifying such signals. The identification of the KOXII factors and their ability to augment the niche functions of MSCs *ex vivo* through the generation of revitalized MSCs (rMSCs) provided a unique platform for a proteomic approach to explore HSPC-niche interactions. By generating a new secretomic dataset characterizing the factors secreted by rMSCs, this study offers insights into the protein factors that mediate MSC-HSPC crosstalk. The presence of interferon signatures in both the transcriptome and secretome of rMSCs suggest that this dataset is especially relevant for HSPC-niche interactions in the context of hematopoietic stress and inflammation.

Additionally, this dataset is not limited to MSC-derived signals supporting HSPCs. Data from our co-culture secretomic screen reveals a large-scale increase in protein secretion by MSCs in response to co-culture with HSPCs. This suggests either that MSC-HSPC crosstalk leads to major changes in the secretory behavior of MSCs, or that the presence of hematopoietic cells drives large scale release of factors previously trapped within the ECM. In the future, this rMSC co-culture system could prove useful in studying HSPC-niche interactions more closely and uncovering the mechanisms through which HSPCs might augment the secretion of niche factors by MSCs.

Furthermore, this study provides evidence supporting a novel role of SEM3A in direct regulation of HSPC proliferation. Previous studies have clearly established SEM3A as a negative regulator of vascular growth, driving endothelial cell apoptosis and slowing vascular regeneration in the bone marrow following total body irradiation and chemoablation (Guttmann-Raviv et al., 2007; Termini et al., 2021). However, these studies have been equivocal on a direct role of SEM3A in regulating HSPCs. Targeted deletion of *Sema3a* with the endothelial cell-specific *Cdh5*-CreER^T2^ had no effect on peripheral blood counts or HSPC populations within the bone marrow at steady state but did accelerate hematopoietic recovery after irradiation, which the authors attributed to accelerated vascular recovery (Termini et al., 2021). Here, we have provided evidence for direct regulation of HSPCs by SEM3A, showing that, in the context of an *ex vivo* culture system which promotes HSC proliferation, SEM3A preserves HSCs by enforcing quiescence. Conversely, we have shown that mice bearing the germline *Sema3a*^K108N^ loss-of-function mutation exhibit HSC and MPP expansion, along with increased phosphorylation of multiple cytokine receptors in HSPCs, suggesting that loss of *Sema3a* leads to dysregulation of HSPC proliferation *in vivo*. Consistent with this idea, Sema3a^K108N^ mutant mice also exhibit accelerated erythrocyte and platelet recovery after 5-FU. As megakaryocyte and erythrocyte-biased multipotent progenitors (MPP^MkE^) were the only MPP population not to be expanded in *Sema3a*^KN/KN^ mice, these results may indicate a relative survival advantage of MPP^MkE^ in *Sema3a*^KN/KN^ mice in the context of 5-FU treatment. Notably, we do not see an increase in WBC recovery after 5-FU, which has been shown to accompany accelerated vascular recovery in endothelial cell-targeted *Sema3a* deletion, suggesting that this effect is not mediated through changes in vascular recovery (Termini et al., 2021).

This work also provides the first evidence that MSC-derived SEM3A plays a role in the bone marrow microenvironment. While *Sema3a* RNA expression is mostly restricted to the sinusoidal endothelial cell compartment at steady state, 5-FU treatment leads to an upregulation of *Sema3a* expression in MSCs, suggesting that this is part of the response of MSCs to hematopoietic stress. This is in keeping with our finding that rMSCs, where SEM3A is highly upregulated at both the RNA and protein levels, exhibit transcriptomic and secretomic signatures of inflammation. If MSCs constitute a distinct source of SEM3A within the bone marrow, this may also explain why we observe changes in hematopoiesis with a germline *Sema3a*^K108N^ loss-of-function mutant that were not previously seen with endothelial-targeted deletion of *Sema3a*. SEM3A is sensitive to extracellular proteases, and it is possible that proteolytic degradation could restrict its activity to local microenvironments (Adams et al., 1997). This would allow MSC-derived SEM3A to have a distinct regulatory function from endothelial-derived SEM3A.

As the K108N mutation interferes with the plexin binding surface of SEM3A, our results in Sema3A^K108N^ mice also provide indirect evidence for a role of plexin receptors in control of HSPC numbers. SEM3A has been shown to bind multiple plexins, including all class A plexins (Plexin A1-4) and Plexin D1 (Worzfeld and Offermanns, 2014). Of these, *Plxna1*, *Plxna3*, and *Plxnd1* are expressed in HSPCs, alongside NRP1, which is thought to be an obligate co-receptor of SEM3A (Toghani et al., 2025; Worzfeld and Offermanns, 2014). There is also the possibility of alternative tissue-specific receptors of SEM3A, allowing for differential responses by HSPCs and endothelial cells to SEM3A within the bone marrow. For instance, CD72, which is robustly expressed in HSCs (**Fig. S4, D and E**), has been posited as a NRP1-independent receptor for SEM3A in the hematopoietic system (Eiza et al., 2023). Intriguingly, CD72 has been shown to suppress KIT activation and cycling via activation of downstream phosphatases, which could explain the broad decrease in RTK phosphorylation observed in progenitors from *Sema3a*^K108N^ mice (Kataoka et al., 2010). Therefore, additional work is necessary to identify the specific receptors activated by SEM3A in HSPCs, and to further clarify how these receptors mediate the effects of SEM3A.

More broadly, these results add to emerging evidence for the semaphorin family as mediators of HSPC-niche interactions. Namely, myeloid-derived SEM4A has been shown to promote quiescence and self-renewal in myeloid-biased HSCs, serving as a negative feedback loop for myeloid differentiation (Toghani et al., 2025). Additionally, other members of the semaphorin family are robustly expressed in the bone marrow, either by bone marrow stromal cells or by hematopoietic cells themselves. In our view, the semaphorins are a heretofore underappreciated family of signaling factors with robust expression in the bone marrow, and it warrants further investigation to determine what roles other members of this family may play in the HSPC niche.

## Methods

### Mice

Mice were maintained under pathogen-free conditions in a barrier facility in microisolator cages based on a protocol approved by the Institutional Animal Care and Use Committee (IACUC) at Albert Einstein College of Medicine. Prior approval was obtained from the IACUC for all experimental procedures, and this research was conducted in compliance with all relevant ethical regulations. C57BL/6 mice were obtained from Taconic Biosciences or Jackson Laboratory. *Sema3a*^K108N^ mice were obtained from the Jackson Laboratory (C3;B6-*Sema3a*^m808Ddg^/J). All mice were analyzed at 8 – 12 weeks of age, and mice of both sexes were used, unless otherwise noted. For 5-FU treatment, mice were injected intraperitoneally with 180mg/kg of 5-FU (Sigma-Aldrich). For analysis of peripheral blood, mice were bled retro-orbitally under isofluorane anesthesia and complete blood counts were obtained using an ADVIA 2120i hematology system (Siemens Healthineers).

### Sample preparation for flow cytometry analysis and cell sorting

For HSPC flow cytometry analysis of bone marrow from *Sema3a*^K108N^ mice, femurs were flushed and dissociated in PEB (2 mM EDTA and 0.5% BSA in PBS) using a 21-gauge needle. The absolute number for each population per femur was calculated based of the total number of nucleated cells flushed from one femur, determined using an ADVIA 2120i hematology system (Siemens Healthineers). For isolation of bone marrow stromal cells, marrow plugs were flushed and digested for 30min at 37°C in 1 mg/mL of dispase II (Gibco) and 2 mg/mL of collagenase IV (Gibco) in HBSS (Corning). After RBC lysis with Ammonium-Chloride-Potassium (ACK) buffer, cells were filtered, stained for cell surface markers, resuspended in PEB with 4′,6-diamino-2-phenylindole (DAPI; Sigma) for dead cell discrimination, and analyzed. Dead cell, doublets, and debris were excluded by forward scatter (FSC), side scatter (SSC), and DAPI. Intracellular Ki67 and DAPI staining was performed as previously described (Jalbert and Pietras, 2018). Briefly, after staining for surface markers, cells were fixed and permeabilized using the Cytofix/Cytoperm Fixation/Permeabilization Kit (BD) following manufacturer instructions. Cells were stained with α-Ki67-FITC for 30min at room temperature, washed, and resuspended in PBS + DAPI for acquisition (25ng/mL; Sigma-Aldrich). Flow cytometry was performed using a LSRII flow cytometer (BD Biosciences) or a Cytek Aurora spectral flow cytometer (Cytek Biosciences). Flow cytometry-assisted cell sorting was performed on a FACSAria II cell sorter (BD Biosciences). Flow cytometric data was analyzed using FlowJo 10 (FlowJo).

### Ex vivo HSPC culture

For culture of Lin-depleted cells in conditioned media, lineage depletion of murine bone marrow was performed as previously described (Nakahara et al., 2019). StemSpan serum free expansion media (STEMCELL Technologies), supplemented with 10% KnockOut Serum Replacement (Gibco) and 100U/mL penicillin/streptomycin (Gibco), was conditioned over rMSCs or cMSCs for 24h. For trypsin pre-treatment, conditioned media was treated with 100μg/mL of trypsin (Sigma-Aldrich) overnight, which was inactivated with 500μg/mL of trypsin inhibitor (Sigma-Aldrich). Trypsin-treated or non-treated conditioned media was diluted 1:1 with non-conditioned media and supplemented with SCF (10ng/mL; Peprotech) and TPO (20ng/mL; Peprotech) before being used to culture Lin-depleted bone marrow for 6d at 37°C, 20% O_2_, and 5% CO_2_.

For analysis of *ex vivo* expansion with recombinant SEM3A, a polyvinyl alcohol- (PVA-) based HSPC culture system was adapted from exiting protocols (Wilkinson et al., 2019; Wilkinson et al., 2020). Bones were collected, flushed, Lin-depleted, and stained for sorting. BSA was excluded from all buffers used in this process. 10μg/mL of recombinant SEM3A-Fc (Macromolecular Therapeutics Development Facility, Albert Einstein College of Medicine) or IgG2a control (R&D Systems) was added to culture media, consisting of Ham’s F-12 (Gibco) supplemented with 1mg/mL PVA (87–90%-hydrolyzed; Sigma), L-glutamine (Gibco), insulin-transferrin-selenium-ethanolamine (ITS-X; Gibco), 10mM HEPES (Gibco), and 100U/mL penicillin/streptomycin (Gibco). 2,000 LSKs were sorted directly into culture media in flat-bottom 96-well plates pre-coated with 1μg/well of fibronectin (Sigma-Aldrich) for 2h at room temperature immediately prior to cell sorting. Cells were cultured for 4 days at 37°C, 20% O_2_, and 5% CO_2_. After culture, cells were lifted by pipetting, stained for surface markers, spiked with CountBright beads absolute counting beads (ThermoFisher Scientific) to allow for absolute quantification, and analyzed by flow cytometry. For analysis of cell cycle occupancy of *ex vivo* expanded cells, 10,000 LSKs were sorted as above, and cultured for 2 days at 37°C before being analyzed.

### Generation of recombinant SEM3A-Fc

Residues N21-K747 of murine *Sem3a* cDNA (Horizon Discovery Biosciences) were PCR amplified and cloned into a derivative of pcDNA3.3 under the human β-2 microglobulin signal peptide, with a carboxy murine IgG2a Fc. The expression construct was transfected into FreeStyleTM 293-F cells (Gibco) using polyethylenimine (25kDa, linear, Polysciences Inc.), and a stable transfected pool was selected using 800µg/ml G418. Valproic acid was added to 3mM, and culture supernatant harvested by centrifugation 5 days later. Protein was affinity purified (mAbSelectSure, Cytiva), eluted with 0.1M arginine, pH 3.0 and neutralized immediately with 1/10th volume 1M Tris, pH 9, and further purified through gel filtration (Superdex S200 26/60, GE) in PBS. Endotoxin level was determined as <1EU/mg by LAL assay (QCL, Lonza).

### RNA isolation and qPCR

After sorting cells directly in lysis buffer, RNA was purified using the Dynabeads RNA Purification Kit (Invitrogen) and reverse-transcribed using the RNA to cDNA EcoDry Premix (TaKaRa Bio), according to manufacturers’ instructions. qPCR was performed with LightCycler 480 SYBR Green I Master Mix (Roche) on a QuantStudio 6 Real-Time PCR System (Applied Biosystems). Primer sequences are provided in **Table S4**. The average cycle threshold number (Ct) across technical triplicates was used to calculate mRNA abundance relative to *Gapdh* (2^-ΔΔCt^).

### Analysis of SEM3A by ELISA

To collect bone marrow extracellular fluid, femurs were flushed with 1mL of PBS and centrifuged to pellet cells. The supernatants were collected and stored at −80°C. SEM3A protein was quantified using a SEM3A ELISA kit (Biomatik) according to the manufacturer’s instructions. For cell culture supernatants, undiluted cell-free supernatants were collected and stored at −80°C, and SEM3A protein was quantified using a SEM3A ELISA kit (LSBio) according to the manufacturer’s instructions.

### Phospho-protein analysis

KIT^+^ BMNCs were enriched from the bone marrow of *Sema3a*^K108N^ mice using α-CD117-bound magnetic beads (Miltenyi Biotec) according to manufacturer’s instructions. For each replicate, 2×10^7^ cells pooled from two mice were used. Receptor phosphorylation was quantified using the Mouse RTK Phosphorylation Array C1 (RayBiotech) according to manufacturer’s instructions.

### Proteomic sample preparation for pSILAC

1.5×10^5^ cMSCs or rMSCs were seeded in 10cm culture-treated plates in their normal growth media, consisting of αMEM (Corning), supplemented with 20% FBS (Gibco), L-glutamine (Gibco), 100U/mL penicillin/streptomycin (Gibco), and 5 ng/mL βFGF. After 24h, media was switched to FBS- and βFGF-free growth media. SILAC-compatible αMEM was prepared by supplementing SILAC-compatible MEM (Athena Enzyme Systems) with non-essential amino acids (Gibco), 1mM sodium pyruvate (Gibco), 200ng/mL lipoic acid (Sigma), 50μg/mL L-ascorbic acid (Sigma), 100ng/mL d-biotin (Sigma), 1.36 μg/mL vitamin B12 (Sigma). After 24h of serum starvation, media was switched to SILAC-compatible αMEM without L-arginine or L-proline to deplete cellular stores of both amino acids. After 20 minutes of amino acid depletion, media was switched to SILAC-compatible αMEM supplemented with either ^13^C^15^N-labelled L-arginine and L-proline (heavy-labelled condition) or ^13^C-labelled L-arginine and d_4_-labelled L-proline (intermediate-labelled condition) (Silantes). One replicate was label swapped. After 12 hours, conditioned media was collected, centrifuged, filtered, and frozen.

Protein concentration in serum samples was measured with bicinchoninic acid assay according to the manufacturer’s instructions (Thermo Fisher Scientific). 50 μg of protein was diluted in a total volume of 100 μl of 50 mM ammonium bicarbonate (pH 8) and heated for 15 min at 95°C for denaturation. Disulfide bonds were reduced and alkylated for 5□min at 50□°C with 10□mmol·L^−1^ DTT and for 25□minutes at 45□°C with 40□mmol·L^−1^ IAA in the dark, and the reaction was quenched with 20□mmol·L^−1^ DTT at RT. Digestion with Trypsin/Lys-C (Promega) was performed in a protease-to-protein ratio of 1:50 (w/w) overnight at 37°C with shaking at 700 rpm. After digestion, trifluoroacetic acid was added (pH < 2) to a final concentration of 1%. Mass spectrometry (MS) injection-ready samples were stored at −20°C.

### LC-MS/MS acquisition and data analysis for pSILAC

For LC-MS/MS analysis, peptides were separated using an Easy NanoLC 1200 fitted with a trapping (Acclaim PepMap C18, 5□μm, 100□Å, 100□μm□×□2□cm) and an analytical column (Waters nanoEase MZ Peptide BEH C18, 130□Å, 1.7□µm, 75□µm□×□25□cm). Solvent A was 0.1% (*v*/*v*) formic acid (FA) in ddH_2_O, and solvent B was 80% ACN and 0.1% (*v*/*v*) FA in ddH_2_O. Samples were loaded onto the trapping column with a constant flow of solvent A at a maximum pressure of 800□bar. Peptides were eluted at a constant flow of 0.3□μL·min^−1^ and temperature of 55□°C and maintained using a HotSleevePlus column oven (Analytical Sales and Services). During elution, the percentage of solvent B was increased linearly from 3 to 8% in 4□min, from 8% to 10% in 2□min, from 10% to 32% in 68□min, from 32% to 50% in 12□min, and from 50% to 100% in 1□min. Finally, the gradient was finished with 7□min in solvent B, followed by 11□min at 97% solvent A.

Peptides were introduced into mass spectrometers via a Pico-Tip Emitter 360□μm OD□×□20□μm ID; 10□μm tip (New Objective). The capillary temperature was set at 275□°C. Samples were analyzed on a Q-Exactive HF Orbitrap mass spectrometer (Thermo Fisher Scientific) with a spray voltage of 2□kV. Full-scan MS spectra with a mass range of m/z 350 to 1 500 were acquired with the Orbitrap at a resolution of 60 000 FWHM. The filling time was set to a maximum of 50□ms with an automatic gain control target of 3□×□106□ions. The top 20 most abundant ions per full scan were selected for MS2 acquisition. Dynamic exclusion was set for 25□s. Isotopes, unassigned charges, and charges >7 were excluded. For MS2 scans, the normalized collision energy was set to 26, and the resolution was set to 15 000 FWHM with automatic gain control of 1□×□105□ions and a maximum fill time of 50□ms. The isolation window was set to m/z 2, with a fixed first mass of m/z 110.

The MS were processed with MaxQuant (1.6.2.6) using the Andromeda search engine against UniProtKB/Swiss-Prot databases of Mus musculus (2018.07.16/UP000000589_10090.fasta), with the following search settings: digestion enzyme was set to trypsin/P-LysC, with a maximum of two missed cleavages allowed. SILAC parameters were specified as 13C15N-labelled L-arginine and L-proline (heavy-labelled condition) or 13C-labelled L-arginine and d4-labelled L-proline (intermediate-labelled condition). Precursor and product ion tolerances were set at 20 ppm and 0.5 Da, respectively. Carbamidomethylation of cysteine was set as a fixed modification, and oxidation of methionine, sulfation (Y), phosphorylation (STY) were set as variable modifications (max 3 per peptide). The match between run functions was enabled with a time window of 0.5 min and an alignment window of 10 min. A minimum of one unique peptide and a false discovery rate below 0.01 were set for peptide and protein identification. Protein quantification was performed using the label-free quantification (LFQ) algorithm of MaxQuant. MS2 spectra were not required for the LFQ comparison. As a decoy database, reversed sequences of the target database were used. If not stated otherwise, then MaxQuant settings were left as default.

### Sample preparation for unlabeled proteomics

1.5×10^5^ cMSCs or rMSCs were seeded in 10cm culture-treated plates in their normal growth media. After 24h, media was switched to Ham’s F-12 (Gibco) supplemented with L-glutamine (Gibco), 10mM HEPES (Gibco), and 100U/mL penicillin/streptomycin (Gibco). 3×10^5^ KIT^+^ BMNCs were added to co-culture conditions, after being enriched from the bone marrow of C57BL/6 mice using α-CD117-bound magnetic beads (Miltenyi Biotec) according to manufacturer’s instructions. After overnight culture, conditioned media was collected, centrifuged, filtered, and frozen.

Thawed filtered media was treated with 5 mM DTT and 50 mM ammonium bicarbonate (pH = 8) and left on the bench for about 1 hour for disulfide bond reduction. Samples were then alkylated with 20 mM iodoacetamide in the dark for 30 minutes. Afterward, phosphoric acid was added to the sample at a final concentration of 1.2%. Samples were diluted in six volumes of binding buffer (90% methanol and 10 mM ammonium bicarbonate, pH 8.0). After gentle mixing, the protein solution was loaded to an S-trap filter (Protifi) and spun at 500 g for 30 sec. The sample was washed twice with binding buffer. Finally, 1 µg of sequencing grade trypsin (Promega), diluted in 50 mM ammonium bicarbonate, was added into the S-trap filter and samples were digested at 37oC for 18 h. Peptides were eluted in three steps: (i) 40 µl of 50 mM ammonium bicarbonate, (ii) 40 µl of 0.1% TFA and (iii) 40 µl of 60% acetonitrile and 0.1% TFA. The peptide solution was pooled, spun at 1,000 g for 30 sec and dried in a vacuum centrifuge.

Prior to mass spectrometry analysis, samples were desalted using a 96-well plate filter (Orochem) packed with 1 mg of Oasis HLB C-18 resin (Waters). Briefly, the samples were resuspended in 100 µl of 0.1% TFA and loaded onto the HLB resin, which was previously equilibrated using 100 µl of the same buffer. After washing with 100 µl of 0.1% TFA, the samples were eluted with a buffer containing 70 µl of 60% acetonitrile and 0.1% TFA and then dried in a vacuum centrifuge.

### LC-MS/MS acquisition and data analysis for unlabeled proteomics

For LC-MS/MS analysis, samples were resuspended in 10 µl of 0.1% TFA and loaded onto a Dionex RSLC Ultimate 300 (Thermo Scientific), coupled online with an Orbitrap Fusion Lumos (Thermo Scientific). Chromatographic separation was performed with a two-column system, consisting of a C-18 trap cartridge (300 µm ID, 5 mm length) and a picofrit analytical column (75 µm ID, 25 cm length) packed in-house with reversed-phase Repro-Sil Pur C18-AQ 3 µm resin. Peptides were separated using a 90 min gradient from 4-30% buffer B (buffer A: 0.1% formic acid, buffer B: 80% acetonitrile + 0.1% formic acid) at a flow rate of 300 nl/min. The mass spectrometer was set to acquire spectra in a data-dependent acquisition (DDA) mode. Briefly, the full MS scan was set to 300-1200 m/z in the orbitrap with a resolution of 120,000 (at 200 m/z) and an AGC target of 5×10^5^. MS/MS was performed in the ion trap using the top speed mode (2 secs), an AGC target of 1×10^4^ and an HCD collision energy of 35.

Proteome raw files were searched using Proteome Discoverer software (v2.5, Thermo Scientific) using SEQUEST search engine and the SwissProt mouse database (updated April 2023). The search for total proteome included variable modification of N-terminal acetylation, and fixed modification of carbamidomethyl cysteine. Trypsin was specified as the digestive enzyme with up to 2 missed cleavages allowed. Mass tolerance was set to 10 pm for precursor ions and 0.2 Da for product ions. Peptide and protein false discovery rate was set to 1%. Following the search, data was processed as described previously (Aguilan et al., 2020). Briefly, proteins were log2 transformed, normalized by the average value of each sample and missing values were imputed using a normal distribution 2 standard deviations lower than the mean. Statistical regulation was assessed using heteroscedastic T-test (if p-value < 0.05). Data distribution was assumed to be normal but this was not formally tested.

### Statistics and reproducibility

All data is presented as mean ± standard error of mean (SEM), unless otherwise specified. For experiments using mice, including *ex vivo* culture experiments using primary hematopoietic cells, n represents individual mice. For experiments with MSCs, n represents independent cultures. Experiments presented were replicated at least 3 times. Sample sizes were determined by previous experience from our labs (Ames et al., 2023; Hemmati et al., 2019; Nakahara et al., 2019; Wei et al., 2020), and no statistical methods were used to pre-determine sample sizes. Statistical analyses and data visualizations were made with GraphPad Prism 8 (GraphPad Software) and RStudio (Posit Software).

## Data Availability

The raw LC-MS/MS data, process report files, and extracted peptide features for the pSILAC experiments have been deposited in the ProteomeXchange Consortium via the PRIDE partner repository under the accession code **PXD065939**. The raw LC-MS/MS data and annotated spectra for the unlabeled proteomic experiments are available under the accession code **PXD065232**. Code used for analysis and visualization of transcriptomic and proteomic data is available upon reasonable request.

## Supplemental material summary

**Fig. S1** summarizes the results of label-free proteomic characterization of the cMSC and rMSC secretomes in co-culture with KIT^+^ BMNCs, showing an enrichment of intracellular proteins detected in the cMSC conditions. **Fig. S2** summarizes the results of pSILAC-characterization of the cMSC and rMSC secretomes, showing bone- and ECM-related proteins are enriched in the rMSC secretome and confirming SEM3A is enriched in the rMSC secretome. **Fig. S3** shows data related to *ex vivo* culture of HSPCs, demonstrating that recombinant SEM3A-Fc increases cell viability and demonstrating that treatment with a SEM3A neutralizing antibody blocks HSC expansion by SEM3A. **Fig. S4**, **A** shows gating schema for sorting mature bone marrow leukocytes for evaluation of SEM3A expression in the hematopoietic compartment. **Fig. S4**, **B**-**E** show data from published RNAseq datasets evaluating expression of SEM3A and its receptors in the bone marrow. **Fig. S5** shows data from *Sema3a*^K108N^ mice, demonstrating normal peripheral blood parameters and normal KITL and SEM3A expression levels in the bone marrow at steady state, and accelerated platelet recovery post-5-FU. **Table S1-3** show data and statistics from secretomic experiments. **Table S4** lists details of antibodies, primers, and other materials used in experiments.

## Acknowledgements

This study is dedicated to the late Dr. Paul S Frenette, under whose mentorship this work began. His passion for his work, his creativity of thought, and his scientific bravery have had a lasting impact on every field of study he touched, and on everyone who had the good fortune to work with him. He is deeply missed.

We thank members of the Frenette and Gritsman laboratories, past and present, for invaluable discussions and advice. We thank C. Prophete, A. Landeros, G. Amatuni, J. Kazmi, M. Bhuiyan, and Y. Wang for technical assistance and administrative support. We are grateful to M. Maryanovich for her invaluable feedback on the manuscript. We thank L. Tesfa, Y. Zhang, andD. Sun for assistance in cell sorting. This work was supported by R01 grants from the National Institutes of Health (NIH) (DK056638 to PSF and KG, DK130895 to KG), the New York State Department of Health NYSTEM IIRP (C029154 and C029570 to PSF), and seed funds from Albert Einstein College of Medicine (to KG). KG was also supported by the Betty & Sheldon Feinberg Senior Faculty Scholar in Cancer Research fund and by a pilot project fund from the Cancer Dormancy Institute of Montefiore Einstein Comprehensive Cancer Center. DKB received support from the NYSTEM Training Program in Stem Cell Research, Training Program in Cellular and Molecular Biology and Genetics (5T32GM007491) and Medical Scientist Training Program (5T32GM007288) T32 NIH training grants, and was the recipient of a National Heart,

Lung, and Blood Institute (NHLBI) Predoctoral National Research Service Award (F30HL154749). DFC was supported by a DKFZ postdoctoral fellowship. Flow cytometry and FACS were performed with support from the Stem Cell FACS and Xenotransplanation Facility and the Flow Cytometry Core Facility at Albert Einstein College of Medicine, the latter of which is supported by the NIH shared instruments grants S10OD026833 and S10OD032169. The Proteomics Core Facility and the Macromolecular Therapeutics Development Facility of Albert Einstein College of Medicine also provided technical and material support. Research reported in this publication was also supported by the Montefiore Einstein Comprehensive Cancer Center Support Grant of the NIH under award number P30CA013330. The Sidoli lab gratefully acknowledges for funding the Hevolution Foundation (AFAR), the Einstein-Mount Sinai Diabetes center, and the NIH Office of the Director (S10OD030286). The content is solely the responsibility of the authors and does not necessarily represent the official views of the NIH.

Author contributions: DKB contributed to all experiments and was responsible for all data analysis and visualization. SPCM and LST contributed to experiments. DKB, PSF, and KG conceptualized the project, designed experiments, and interpreted results. FN generated the rMSCs used in this study and provided technical expertise in MSC culture and co-culture techniques. DFC, JK, and SS assisted in designing proteomic experiments. DFC and SS contributed to analysis and visualization of proteomic experiments. JK and SS supervised and contributed to interpretation of proteomic experiments. SG designed and generated recombinant SEM3A. LS assisted with designing experiments and interpreting results. PSF and KG supervised all research and were responsible for funding acquisition. DKB, SPCM, and KG wrote and edited the original draft of this manuscript, which was reviewed by all authors.

## Author Notes

Disclosures: P.S.F. served as a consultant for Pfizer, received research funding from Ironwood Pharmaceuticals outside the submitted work, and was a shareholder of Cygnal Therapeutics. K.G. has received research funding from ADC Therapeutics and iOnctura outside the submitted work. All other authors declare no competing interests.

