## Supplementary Data for "A bone marrow stromal secretome screen identifies semaphorin 3A as a regulator of hematopoiesis"

**Supplemental Figures**

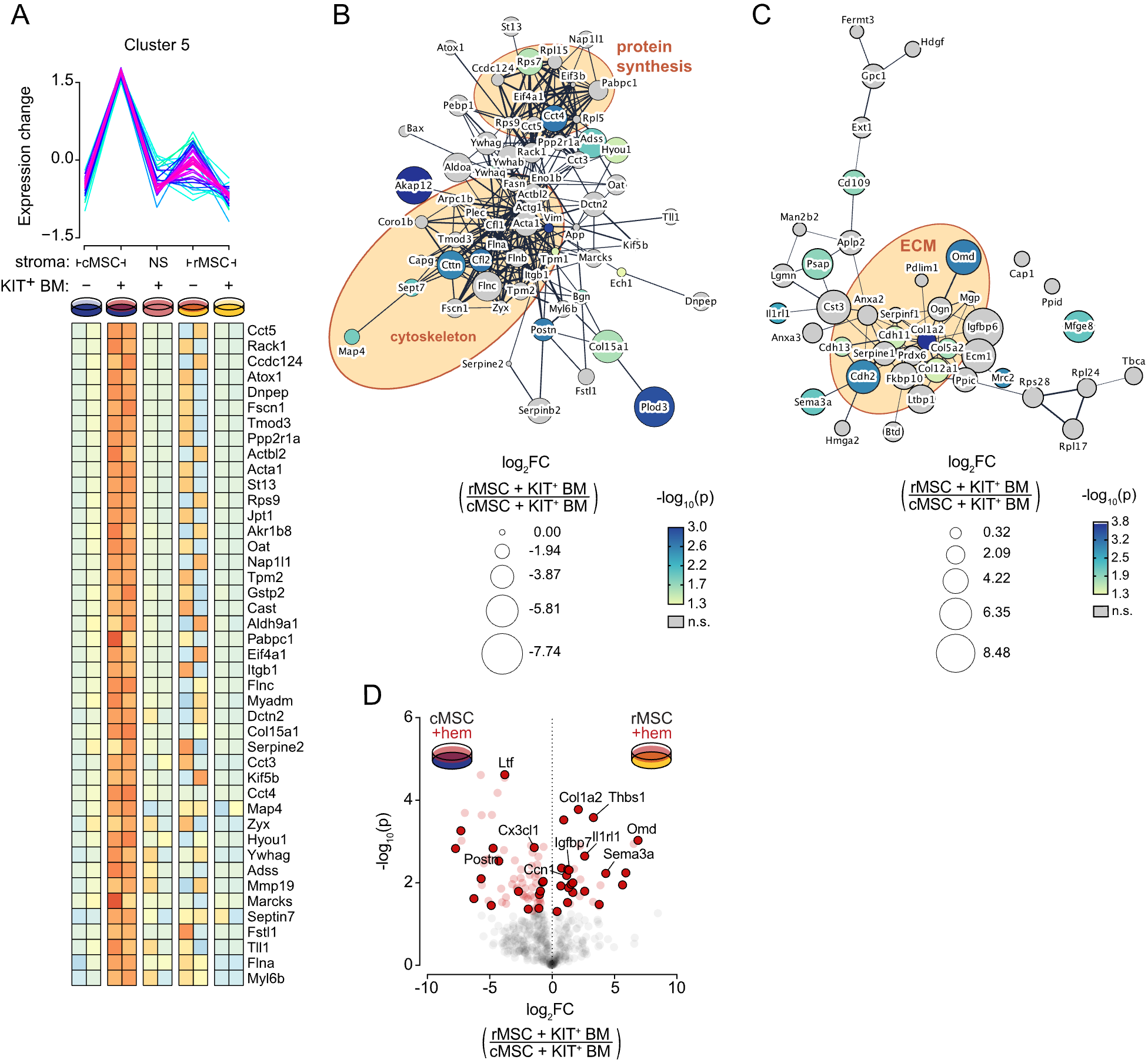

**Figure S1.** (**A**) Pattern of protein detection of cluster 5, demonstrating increased levels in the cMSC secretome in co-culture conditions. Heatmap shows proteins assigned to cluster 1 with membership degree greater than 0.4. (**B**) STRING network of proteins assigned to cluster 5, demonstrating clusters of intracellular machinery detected in media. (**C**) STRING network of proteins assigned to cluster 1, demonstrating a cluster of ECM-related proteins. For **B** & **C**, node size represents log_2_FC of normalized LFQ of proteins in rMSC versus cMSC in co-culture conditions, and node color represents the corresponding p-value. (**D**) Volcano plot showing log_2_FC of normalized LFQ of proteins in rMSC secretome versus cMSC secretome in co-culture conditions, corresponding to **Figure 2D**. Significantly differentially detected proteins (p<0.05) are indicated in red, and extracellular proteins per STRING indicated in dark red.

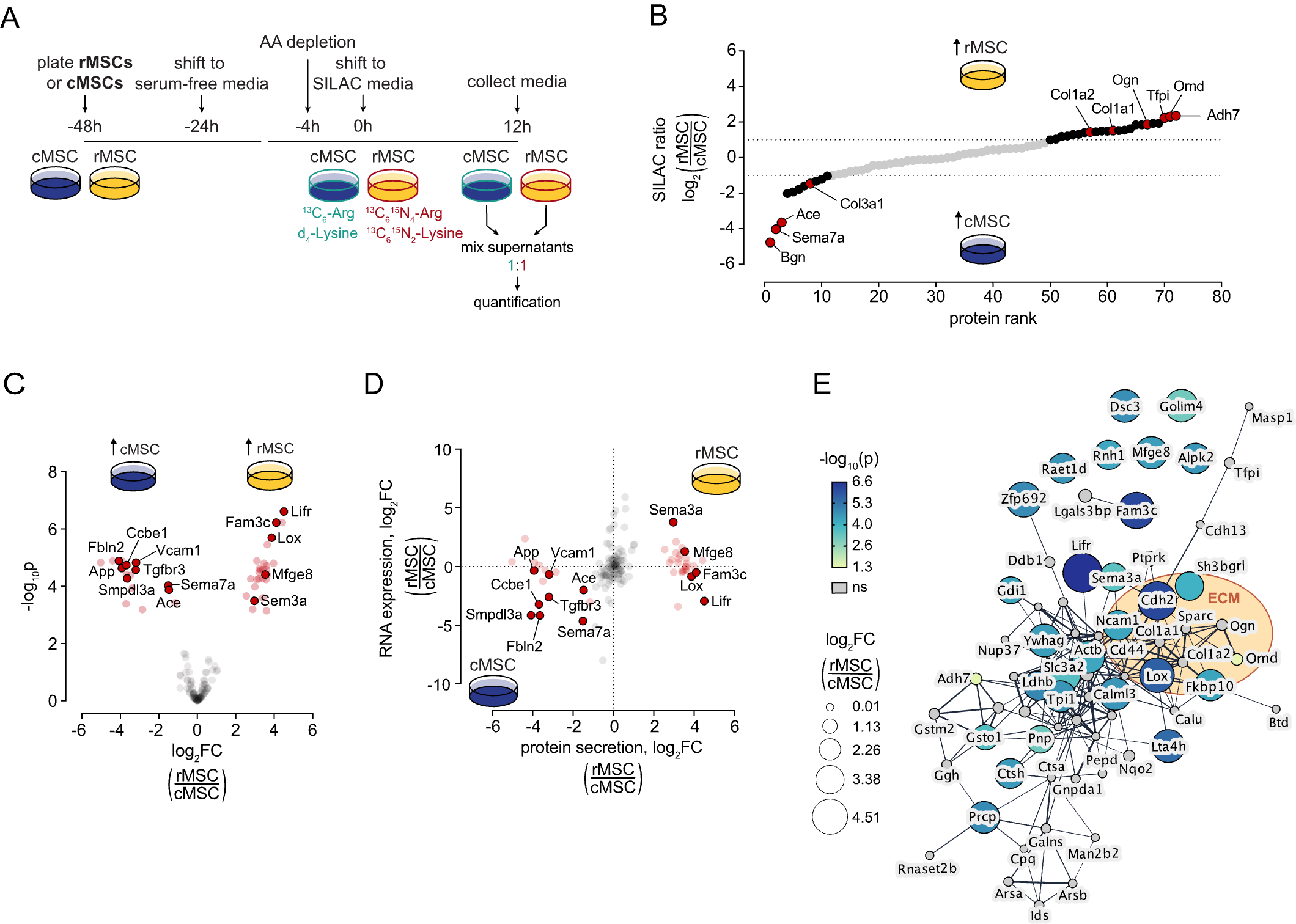

**Figure S2.** (**A**) pSILAC approach for evaluating the rMSC secretome. (**B**) Proteins ranked by SILAC ratio (KOXII/C), with selected proteins labelled. (**C**) Volcano plot showing log_2_FC of normalized LFQ of proteins in rMSC secretome versus cMSC secretome. (**D**) Log fold change (log_2_FC) of normalized label-free quantification (LFQ) of proteins in rMSC secretome versus cMSC secretome plotted against log_2_FC of RPKM of corresponding transcripts, as determined by RNAseq (Nakahara et al., 2019). Significantly differentially detected proteins are indicated in red, with proteins with known secretory functions per UniProt outlined and in dark red. (**E**) STRING network of proteins upregulated in the secretome of rMSCs, demonstrating a cluster of ECM-related proteins. Node size represents log_2_FC of normalized LFQ of proteins in rMSC versus cMSC in co-culture conditions, and node color represents the corresponding p-value.

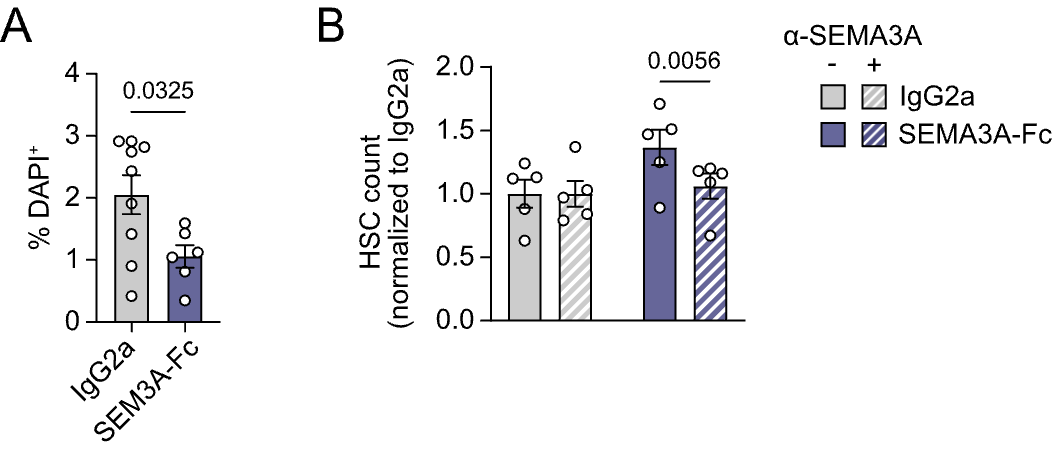

**Figure S3.** (**A**) % DAPI+ cells after 4d of culture of 2x10^3^ LSKs with recombinant Sema3a-Fc or IgG2a isotype control. (**B**) CD150+ EPCR+ HSC numbers after 4d of culture of 2x10^3^ LSKs with recombinant Sema3a-Fc or IgG2a isotype control, with or without an anti-Sema3a blocking antibody.

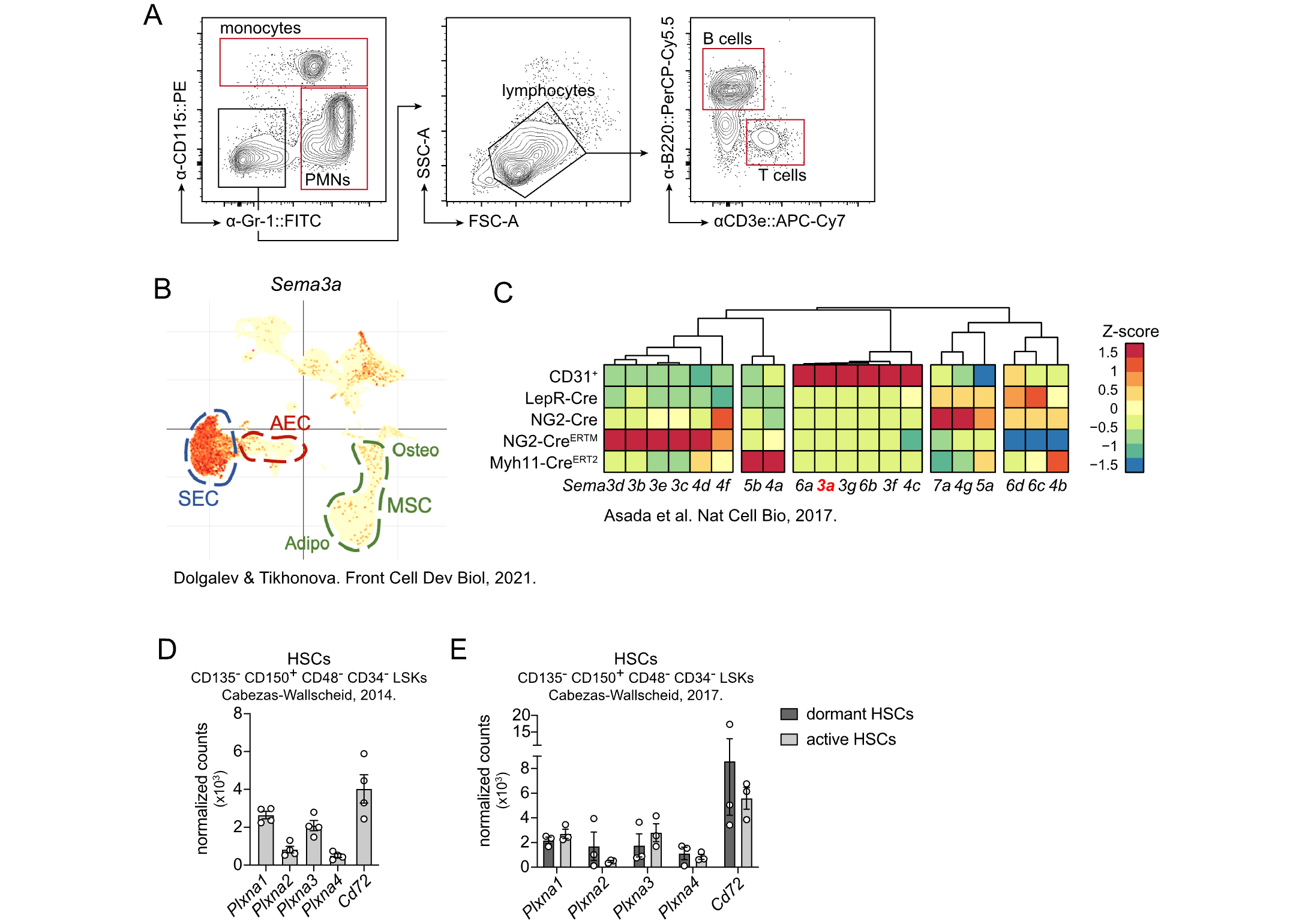

**Figure S4.** (**A**) Sort approach for mature BM leukocytes used for **Figure 4A**. (**B**) Expression of *Sema3a* in murine BM stromal cells, as determined by scRNAseq (Dolgalev and Tikhonova, 2021). (**C**) Expression of semaphorin family members in niche populations *in vivo*, as determined by RNAseq of endothelial cells (CD31+) and Cre-labelled MSC populations, with *Sema3a* showing high expression in endothelial cells (Asada et al., 2017). (**D**) mRNA expression of SEM3A receptors and co-receptors in HSCs, as determined by RNAseq, from (Cabezas-Wallscheid et al., 2014) and (**E**) (Cabezas-Wallscheid et al., 2017).

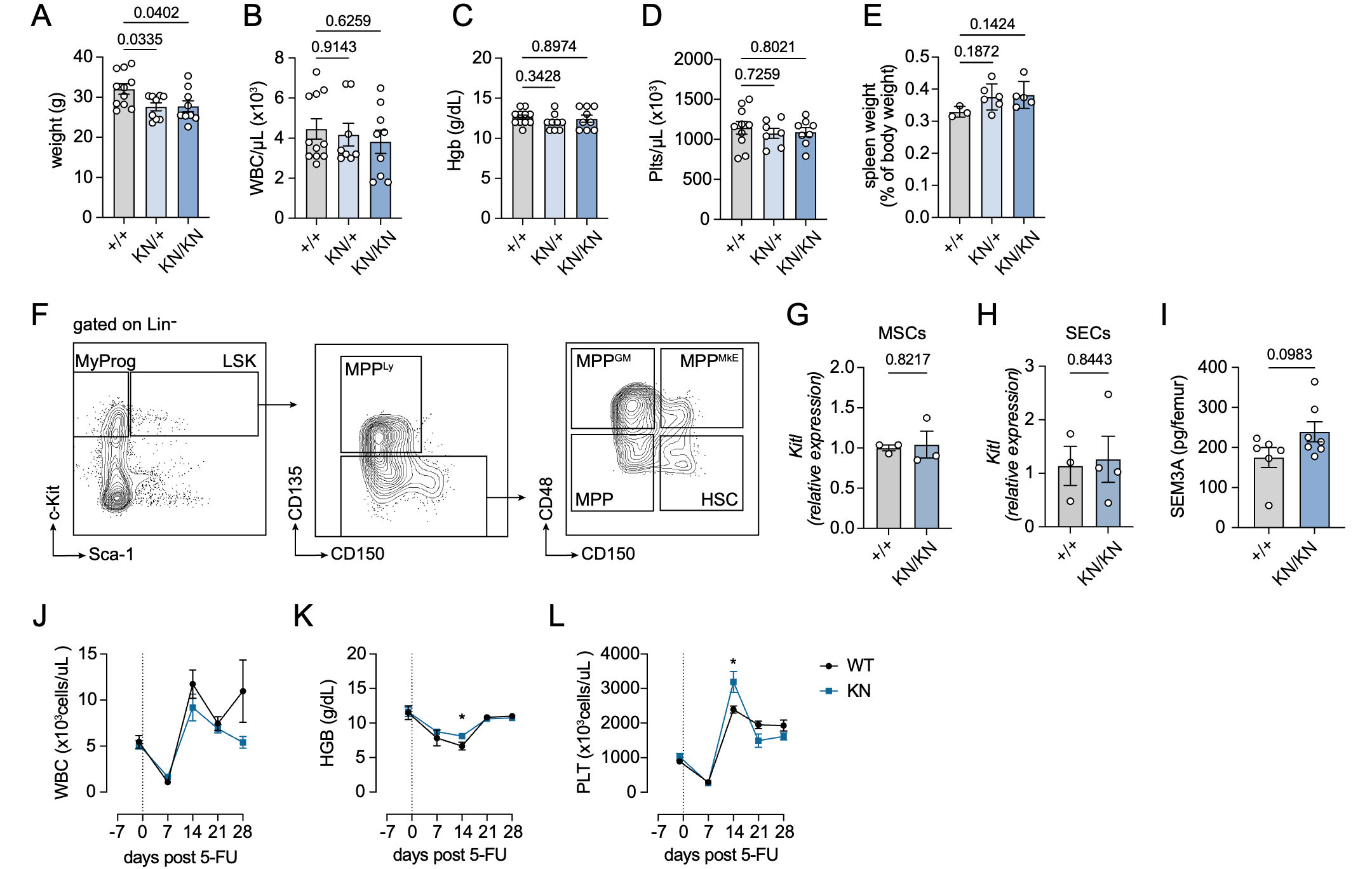

**Figure S5.** (**A**) Body weights of *Sema3a*^+/+^, *Sema3a*^+/K108N^, and *Sema3a*^K108N/K108N^ mice. (**B**) Peripheral blood white blood cell counts (WBC), (**D**) hemoglobin concentration (HGB), and (**C**) platelet counts (Plts) in these mice. (**E**) Spleen weights, as a percentage of total body weight, for *Sema3a*^+/+^, *Sema3a*^+/K108N^, and *Sema3a*^K108N/K108N^ mice. (**F**) Gating scheme for assessing HSPCs in the bone marrow of *Sema3a*^K108N^ mice. (**G**) Expression of *Kitl* in MSCs and (**H**) SECs, as determined by qPCR. (**I**) SEM3A ELISA performed on BMEF of *Sema3a*^+/+^ and *Sema3a*^K108N/K108N^ mice. (**J**) WBC, (**K**) HGB, and (**L**) Plt recovery post-5FU injection. For **A** - **E**, ANOVA with Tukey multiple-comparisons test was used to compare groups. For **G** - **L**, unpaired two-tailed Student’s *t* test was used to compare conditions. No correction for multiple testing was used for **J** - **L**.
